# Biophysical mechanisms of morphogenesis in lizard lungs

**DOI:** 10.1101/2025.09.01.673487

**Authors:** Kaleb Hill, Aaron H. Griffing, Michael A. Palmer, Bezia Lemma, Aria Lupo, Tony Gamble, Natalia A. Shylo, Andrej Košmrlj, Paul A. Trainor, Celeste M. Nelson

## Abstract

The lungs of squamate reptiles (lizards and snakes) are highly diverse, exhibiting single chambers, multiple chambers, transitional forms with two to three chambers, along with a suite of other anatomical features, including finger-like epithelial projections into the body cavity known as diverticulae. During embryonic development of the simple, sac-like lungs of anoles, the epithelium is pushed through the openings of a pulmonary smooth muscle mesh by the forces of luminal fluid pressure. This process of stress ball morphogenesis generates the faveolar epithelium typical of squamate lungs. Here, we compared embryonic lung development in brown anoles, leopard geckos, and veiled chameleons to determine if stress ball morphogenesis is conserved across squamates and to understand the physical processes that generate transitional-chambered lungs with diverticulae. We found that epithelial protrusion through the holes in a pulmonary smooth muscle mesh is conserved across squamates.

Surprisingly, however, we found that luminal inflation is not conserved. Instead, leopard geckos and veiled chameleons appear to generate their faveolae via epithelial folding downstream of epithelial proliferation. We also found experimental and computational evidence suggesting that the transitional chambers and diverticulae of veiled chameleon lungs develop via apical constriction, a process known to be crucial for airway branching in the bird lung. Thus, distinct morphogenetic mechanisms generate epithelial diversity in squamate lungs, which may underpin their species-specific physiological and ecological adaptations.

## 1 INTRODUCTION

The vertebrate lung exhibits strikingly diverse epithelial architectures, including the tree-like airways of mammals, the looped airways of birds, and the corrugated, faveolar sacs of amphibians and non-avian reptiles.^1,2^ The latter span the extremes of epithelial complexity. More complex forms include the multi-chambered and multi-lobed lungs of turtles,^3,4^ the multi-chambered lungs of crocodilians,^3,5^ and the multi-chambered sac-like lungs of some lizards and snakes.^3,4^ Less complex forms include the single-chambered sac-like lungs of most lizards and tuatara.^3,6^ To achieve this variation in epithelial morphology, different vertebrate lineages appear to use distinct morphogenetic mechanisms during early lung development.^7^ The epithelium of the mammalian lung forms branches in response to mechanical constraints from patterns of smooth muscle differentiation.^8–10^ The epithelium of the bird lung folds via apical constriction and later fuses to generate the circuit for unidirectional airflow.^11–14^ The epithelium of the lungs of lizards and clawed frogs expands through openings in a smooth muscle mesh, apparently in response to inflation of the lumen with fluid.^15,16^ This process, known as stress ball morphogenesis,^16^ is required for lung development in anole lizards, but it remains unclear whether stress ball morphogenesis is conserved in other reptiles.

Squamates encompass a lineage of more than 12,000 described species of lizards and snakes with diverse morphologies and physiologies that have facilitated their ecological radiation and adaptation.^17^ Their lungs are similarly diverse and can be categorized into three chamber types: single chamber, transitional chamber, or multi-chamber. All major squamate lineages have species that exhibit single-chambered lungs, which are simple undivided sacs (**Fig. 1A**).^3^ Several lineages within gekkotan, anguimorph, and iguanian lizards exhibit transitional-chambered lungs, which have two or three chambers separated by internal longitudinal septa (**Fig. 1A**).^3,18,19^ Some snakes and anguimorph lizards exhibit multi-chambered lungs with cartilage-reinforced bronchi (**Fig. 1A**).^3,20–22^ In addition, three lineages of squamates exhibit finger-like outpouchings of epithelial tissue known as diverticulae: *Uroplatus* geckos,^23,24^ *Polychrus* lizards,^25^ and chameleons (**Fig. 1A**).^18^ As compared to single-chambered lungs, transitional and multi-chambered lungs are thought to have increased surface area for gas exchange,^26,27^ an ability to store air,^3^ and a morphology that permits unidirectional airflow.^4,19^ The function of diverticulae remains unknown; however, these epithelial outpouchings may increase lung volume in arboreal lineages that have compressed torsos.^18,26^ The morphogenetic mechanisms that build these more complex epithelial structures are also unknown.

**Figure 1.**
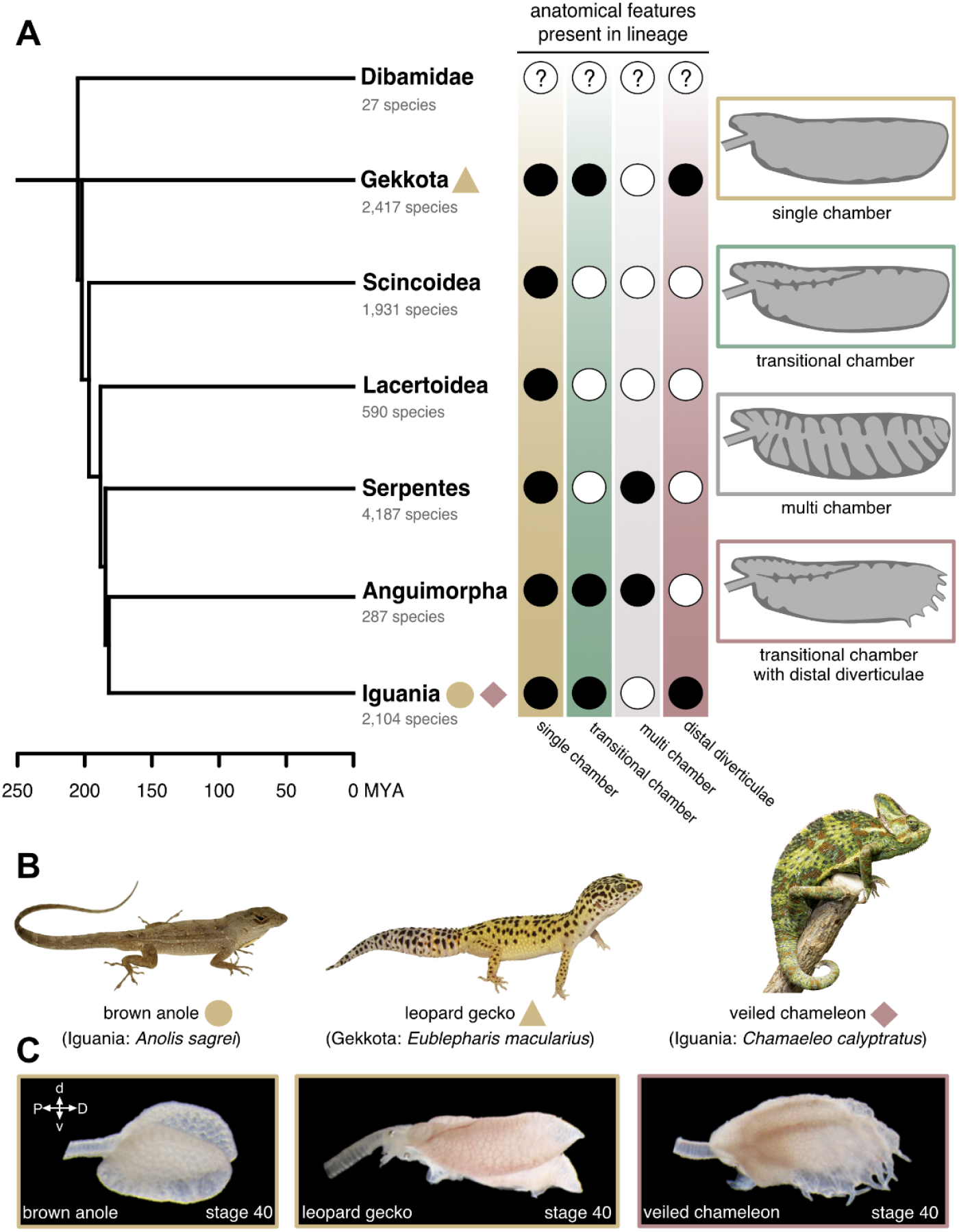
Lung diversity in squamate reptiles. **A**) Major lineages of squamate reptiles in a phylogenetic context and anatomical features present in each lineage. Black and white circles correspond to presence and absence of anatomical features in the lineage, respectively. Anatomical data from Perry (1998). Phylogeny modified from Zheng and Wiens (2016). **B**) Photographs of squamate species investigated in this study. **C**) Lateral view of near-hatching embryonic lungs of species investigated in this study. MYA, millions of years ago; P, proximal; D, distal; d, dorsal; v, ventral.

Here, we investigated embryonic lung development in three species of squamates to identify the extent to which stress ball morphogenesis is conserved and to uncover the physical mechanisms that generate transitional chambers and diverticulae. We found that meshes of pulmonary smooth muscle are conserved across squamates; however, different species generate faveolae via epithelial growth through the mesh, as opposed to luminal pressure pushing the epithelium through the mesh. We found that the individual chambers and diverticulae of the chameleon lung appear to initiate via apical constriction, the process that induces branch formation in the avian lung. Taken together, our results suggest that a variety of morphogenetic mechanisms are used to generate the diverse epithelial morphologies observed in squamate lungs.

## 2 RESULTS

We focused our analysis on three model squamate species: the brown anole (*Anolis sagrei*), leopard gecko (*Eublepharis macularius*), and veiled chameleon (*Chamaeleo calyptratus*; **Fig. 1B**). Both brown anoles and veiled chameleons are iguanian lizards, while leopard geckos are gekkotan lizards (**Fig. 1A**). Because gekkotan lizards, with the possible exception of dibamids, are the sister lineage to all remaining squamates, our taxon sampling scheme spans almost the entire evolutionary history of Squamata.^28^ Before hatching, the lungs of all three species are structurally similar to adult lungs. Both brown anoles and leopard geckos have single-chambered lungs with slight proximal subchambers (**Fig. 1C**).^3,16,29^ Veiled chameleons have lungs with three main chambers and distally situated diverticulae.^18^

Development *in ovo* is variable in squamates and sensitive to temperature. When incubated at 27±1ºC, brown anoles hatch after ∼27 days,^30^ leopard geckos hatch after ∼52 days,^31^ and veiled chameleons hatch after ∼200 days.^32^ We therefore staged embryos using morphological criteria.^30,31,33–35^ Lungs were dissected from embryos between stages 32 and 38 and labeled for markers of epithelium and smooth muscle using immunofluorescence. At stage 32, the brown anole lung begins as a simple wishbone-shaped epithelium without pulmonary smooth muscle (**Fig. 2A**). During stage 33, the lung begins to inflate and, soon after, smooth muscle begins differentiating and surrounding the epithelium (**Fig. 2B, C**). At stage 34, the pulmonary smooth muscle forms into a meshwork and the lumen of the lung further inflates with fluid, decreasing the aspect ratio of each simple sac (**Fig. 2D, E**). A proximal subchamber is visible during this stage (**Fig. 2D**). From stage 34 onward, inflation of the lumen pushes the epithelium through the holes in the smooth muscle mesh (**Fig. 2D–H**), generating the initial morphology of the faveolae.^16^

**Figure 2.**
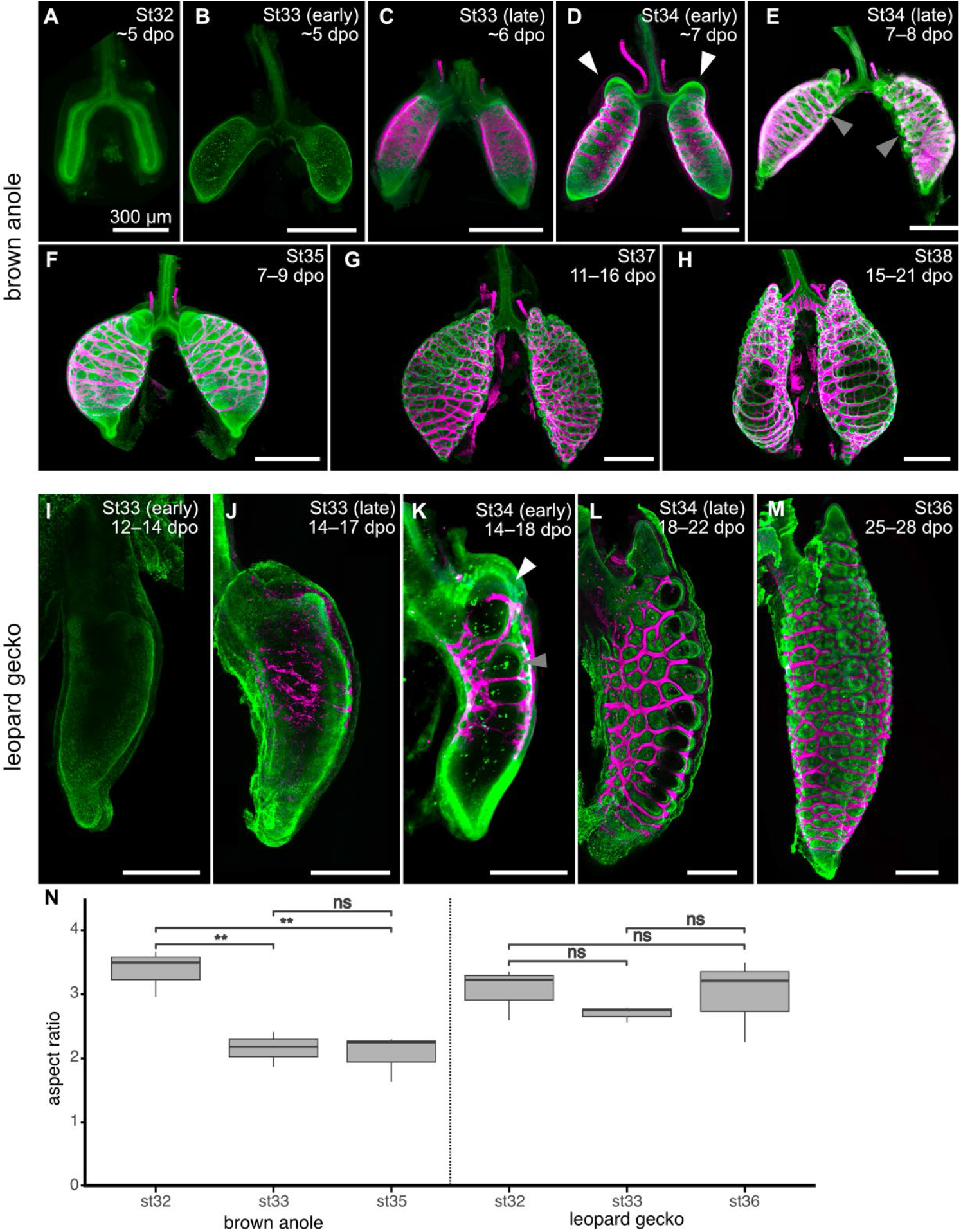
Developmental staging of brown anole and leopard gecko lungs. Images show staining for F-actin (green; **A**) or immunofluorescence for E-cadherin (green; **B–M**) and alpha smooth muscle actin (aSMA; magenta). dpo, days postoviposition; St, developmental stage. Scale bars, 300 µm. **N**) Aspect ratios of developing lungs of brown anoles and leopard geckos. White arrows indicate proximal sub-chamber of both species. Gray arrows indicate epithelium pushing through the smooth muscle mesh. Asterisks and ns denote significant and non-significant differences, respectively, in pairwise *t*-test. Brown anole st32 vs st33 (*p*=0.005), st32 vs st35 (*p*=0.006), st33 vs st35 (*p*=0.375). Leopard gecko st32 vs st33 (*p*=0.141), st32 vs st35 (*p*=0.440), st33 vs st35 (*p*=0.267).

We found that development of the lung of the leopard gecko follows a slightly different series of morphogenetic events. At stage 33, the epithelium is a simple wishbone-shaped sac, similarly absent of pulmonary smooth muscle (**Fig. 2I**). Between stages 33 and 34, pulmonary smooth muscle differentiates and forms into a meshwork (**Fig. 2J, K**). During late stage 34, the epithelium has begun to protrude through holes in the pulmonary smooth muscle mesh (**Fig. 2L**), and a proximal subchamber is similarly visible. By stage 36, the mesh has become more refined and the initial morphology of the faveolae are established (**Fig. 2M**). However, we noticed that the epithelium of the leopard gecko lung does not appear to inflate during these time points. To confirm this qualitative observation, we measured the aspect ratios of brown anole and leopard gecko lungs before the appearance of smooth muscle and after deformation of the epithelium. These measurements confirmed that the lung of the brown anole inflates during these stages (i.e. decreases in aspect ratio between stages) but the lung of the leopard gecko does not (**Fig. 2N**).

We found that development of the transitional-chambered lung of the veiled chameleon is distinct from that of both the brown anole and leopard gecko. At stage 32, the veiled chameleon lung exhibits a wish-bone shaped epithelium with proximally-situated buds that will become the ventral subbronchi (**Fig. 3A**), with no visible pulmonary smooth muscle. At early stage 33, the ventral subbronchus has elongated and the emerging buds of the dorsal subbronchus are visible (**Fig. 3B**). At late stage 33, pulmonary smooth muscle has begun to differentiate around the epithelium of the main bronchus and the most proximal portion of the ventral subbronchus (**Fig. 3C**). At this stage, an additional sub-chamber is visible proximal to the ventral subbronchus (**Fig. 3C**). At stage 34, both subbronchi and the proximal sub-chamber have elongated (**Fig. 3D, E**). Smooth muscle has begun to differentiate distally around the main bronchus and wrap around the ventral subbronchus (**Fig. 3D, E**). At stage 35, the subbronchi have elongated further while smooth muscle has continued to wrap in a distal direction around the subbronchi and main bronchus (**Fig. 3F, G**). At late stage 35, smooth muscle has formed a meshwork around most of the epithelium, which has started to protrude through the holes in the mesh (**Fig. 3H**). From stage 36 onward, the epithelial protrusions become more prominent and the lung appears to inflate (**Fig. 3I–L**).

**Figure 3.**
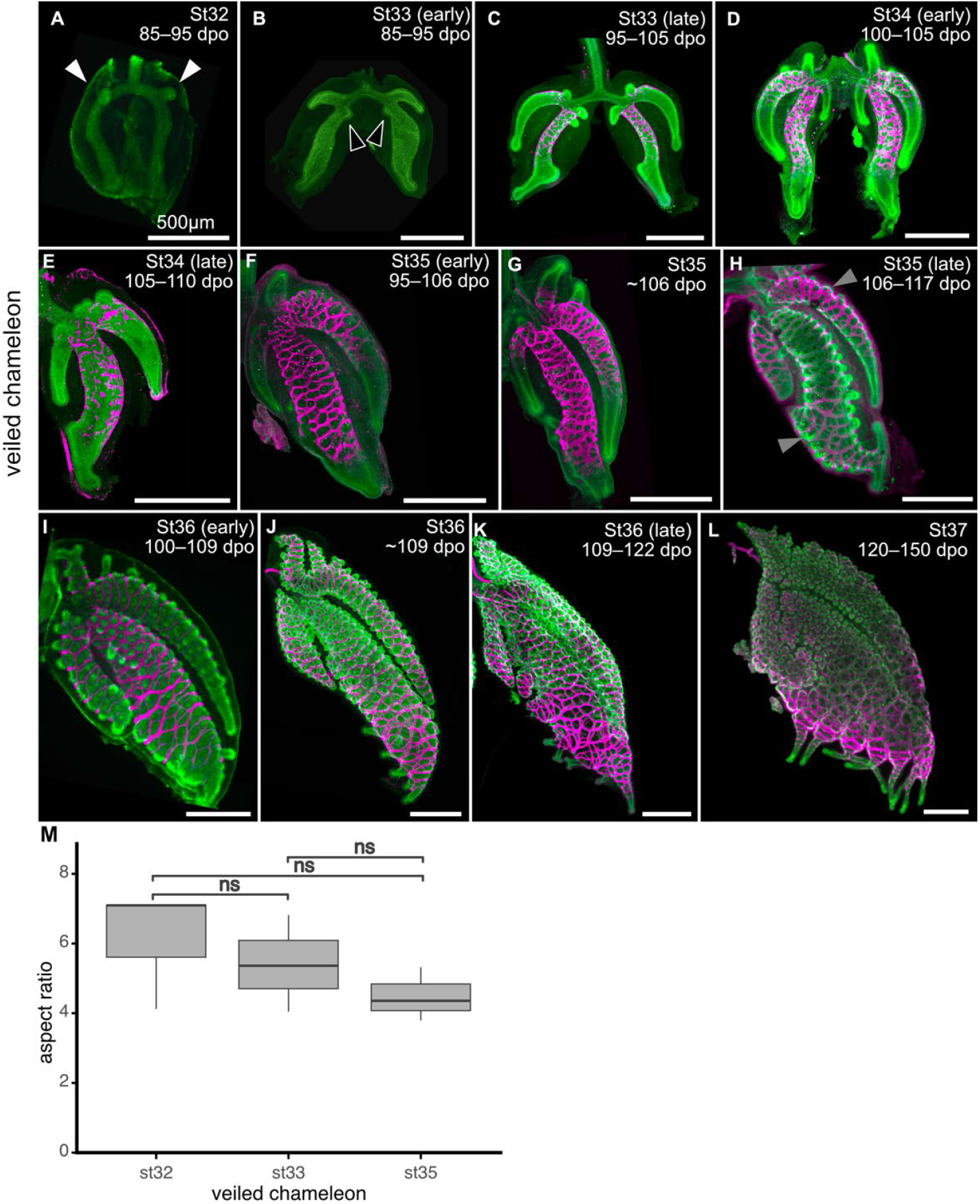
Developmental staging of veiled chameleon lungs. Images show immunofluorescence for aSMA (magenta) and either epithelial cytokeratin (**A, I, L**) or E-cadherin (**B–H, J–K**). dpo, days postoviposition; St, developmental stage, Scale bars, 500 µm. **M**) Aspect ratios of developing lungs of veiled chameleons. White and black arrows indicate emergence of the ventral subbronchus and dorsal subbronchus, respectively. Gray arrows indicate epithelium pushing through the smooth muscle mesh. Asterisks and ns denote significant and non-significant differences, respectively, in pairwise *t*-test. st32 vs st33 (*p*=0.306), st32 vs st35 (*p*=0.117), st33 vs st35 (*p*=0.195).

Nonetheless, the aspect ratio of the veiled chameleon lung remains constant during the stages of development in which the epithelium pushes through the smooth muscle mesh (**Fig. 3M**).

In addition to being transitional-chambered, the veiled chameleon lung differs from those of the brown anole and leopard gecko by having diverticulae, which first appear at stage 36 (**Fig. 4A**). At this stage, smooth muscle wraps around the base of each diverticulum (**Fig. 4A**). At late stage 36, some, but not all, diverticulae have begun to bifurcate (**Fig. 4B**). By stage 37, the diverticulae are long, finger-like structures that resemble the diverticulae within the lungs of adult veiled chameleons (**Fig. 4C, D**). These epithelial structures are more similar in morphology to the branched airways of the mammalian and avian lungs than to faveolae. In the mouse lung, epithelial branches are sculpted by patterns of smooth muscle differentiation,^8–10^ which specifies sites of bifurcation prior to changes in epithelial shape. The epithelium of the diverticulae, however, appears to bifurcate before smooth muscle differentiates at the bifurcation site (**Fig. 4A, B**). In the bird lung, epithelial branches form by active actomyosin-induced contraction of the epithelium itself, in a process known as apical constriction.^11,12^ To determine whether apical constriction might play a role in morphogenesis of the veiled chameleon lung, we conducted immunofluorescence analysis for phosphorylated myosin light chain (pMLC). We observed increased staining intensity for pMLC at the apical surface of the epithelium at the distal tips of the subbronchi and main bronchi from stages 33 to 35 (**Fig. 4E–G**). We also observed a striking enrichment in pMLC at the apical surface of the epithelium in diverticulae (**Fig. 4H**). These data suggest that apical constriction may play a role in initiation of the epithelial folding that constructs both the subbronchi as well as diverticulae in the veiled chameleon lung.

**Figure 4.**
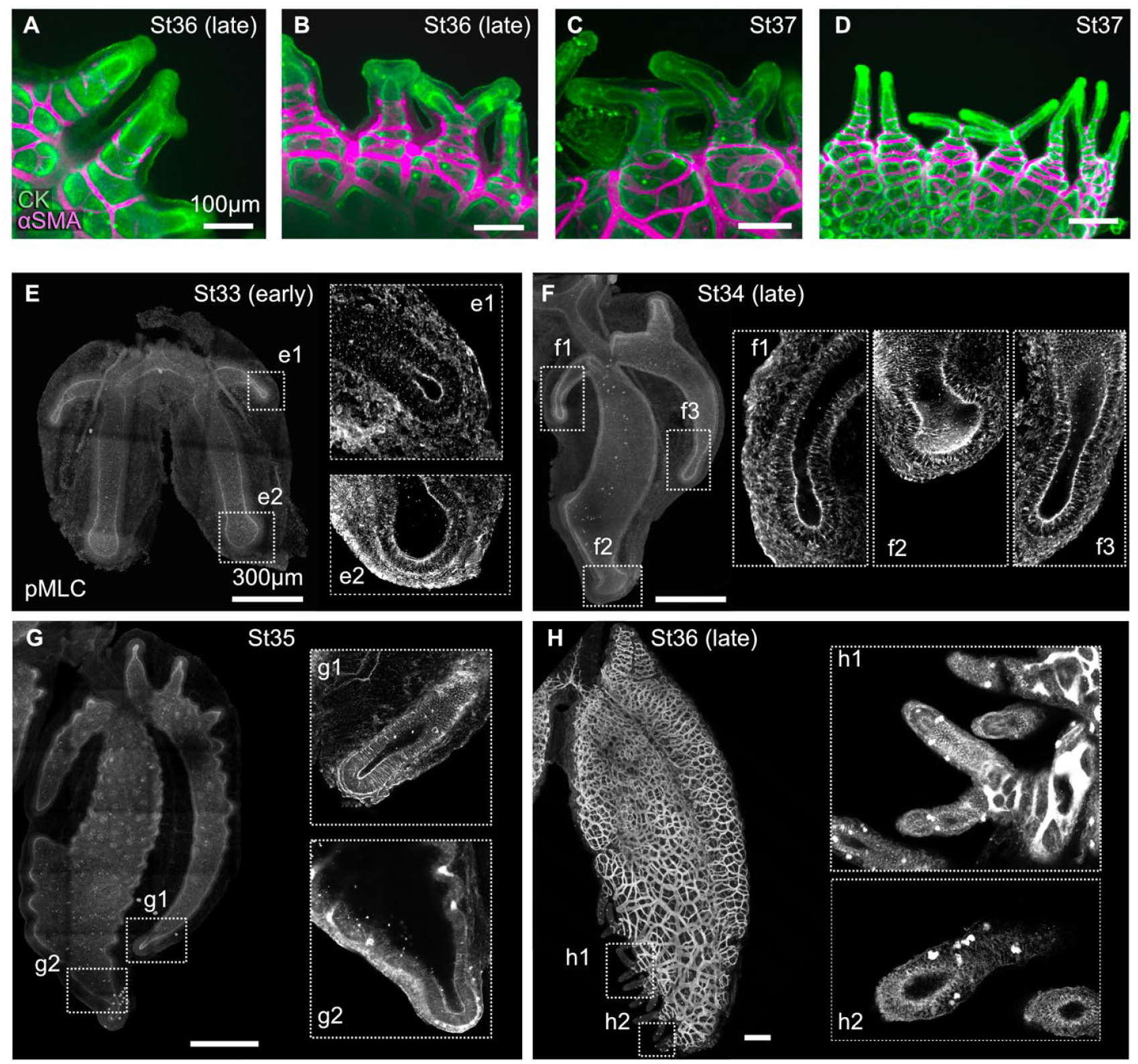
Analysis of smooth muscle coverage and apical constriction in diverticulae and subbronchi of veiled chameleon lungs. **A–D**) Images showing immunofluorescence for cytokeratin (CK) and αSMA in the veiled chameleon lung diverticulae. **E–H**) Images showing immunofluorescence for phosphorylated myosin light chain (pMLC). Scale bars (A–D), 100 µm. Scale bars (E–H), 300 µm.

After the avian airway epithelium changes shape to form a branch, cells at the tip begin to proliferate at a higher rate than the adjacent epithelium, which promotes branch elongation.^11,13^ To determine whether a similar spatial pattern of proliferation accompanies morphogenesis of the subbronchi and diverticulae in the chameleon lung, we isolated lungs at different stages of development and monitored incorporation of the thymidine analog, EdU. From late stage 33 onward, EdU incorporation was substantially higher in epithelial cells located at the growing tips of subbronchi and the proximal sub-chamber than adjacent regions of the lung (**Fig. 5A–F**). Similarly, at stage 37, EdU incorporation was higher in the tips of the diverticulae (**Fig. 5G**). We also found high EdU incorporation in the developing proximal sub-chambers of brown anole (**Fig. 5H**) and leopard gecko (**Fig. 5I**). In the faveolae of both the veiled chameleon and leopard gecko, we observed EdU incorporation in the epithelium as well as the mesenchyme (**Fig. 5J-L**). In contrast, we observed EdU incorporation primarily in the mesenchyme surrounding faveolar epithelium of the brown anole (**Fig. 5M**), consistent with previous observations.^16^ Overall, our data suggest that the faveolar epithelia of leopard geckos and veiled chameleons are highly proliferative. In addition, veiled chameleons exhibit high levels of proliferation in the elongating epithelium of the subbronchi and diverticulae.

**Figure 5.**
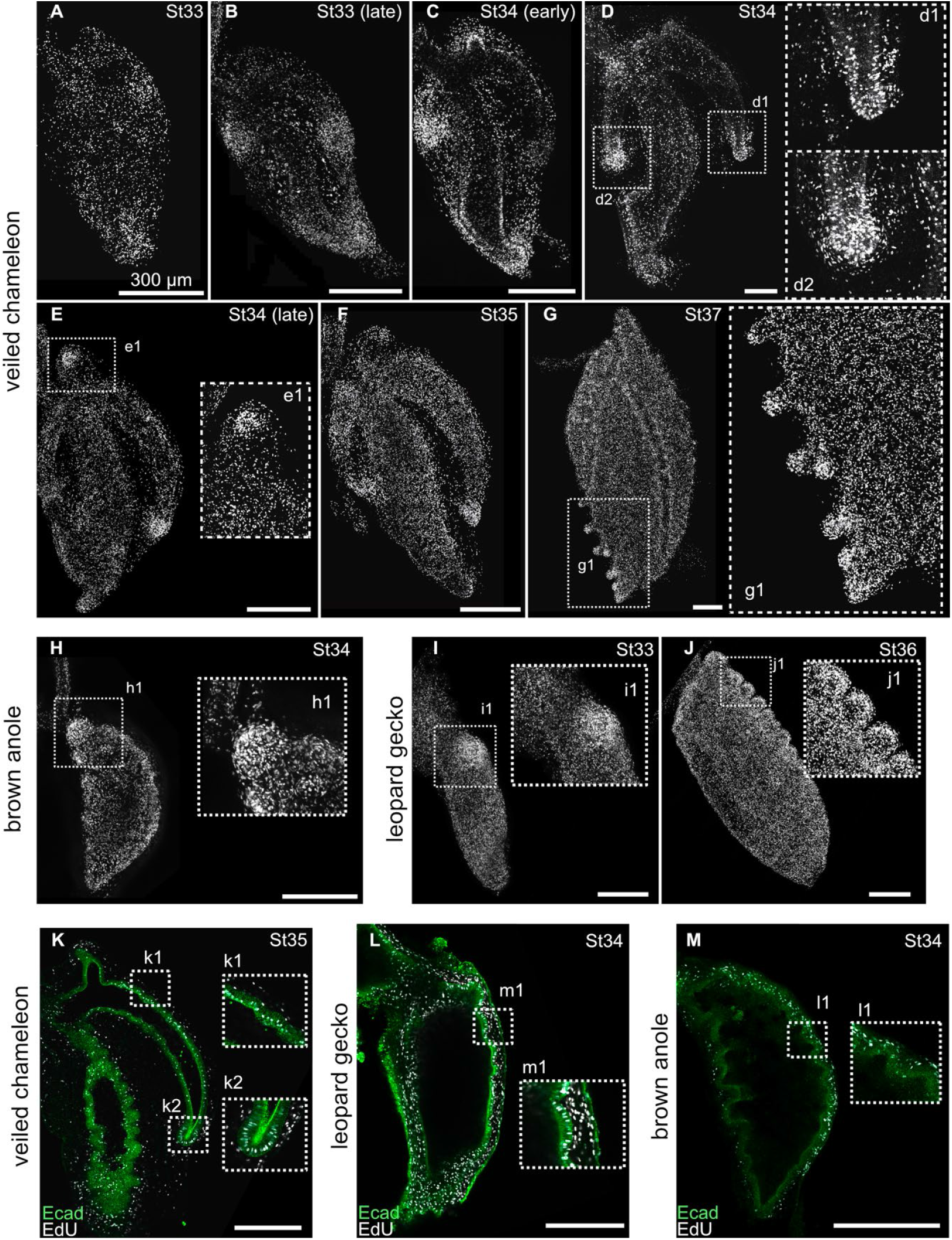
Patterns of cell proliferation in the developing lungs of lizards. **A–G**) Images showing EdU incorporation in the veiled chameleon lung. **H**) Image showing EdU incorporation in the brown anole lung. **I–J**) Images showing EdU incorporation in the leopard gecko lung. **K–M**). Z-slice images showing immunofluorescence for E-cadherin (green) and fluorescent EdU incorporation (cyan) in the veiled chameleon (**K**), leopard gecko (**L**), and brown anole (**M**). Scale bars, 300 µm.

During development of the anole lung, the volume of the fluid within the lumen of the organ increases between stages 32 and 33 (a span of 24 hours) and the resulting pressure is sufficient to expand the epithelium and push it through the holes in the smooth muscle mesh.^16^ That neither the leopard gecko (**Fig. 2N**) nor the veiled chameleon (**Fig. 3M**) show an increase in aspect ratio during faveolar morphogenesis suggests that this process might occur independently of luminal pressurization in these species. Furthermore, the leopard gecko and veiled chameleon exhibit substantial proliferation in the regions of epithelium that fold into faveolae (**Fig. 5K, L**). We hypothesized that the growth from this elevated proliferation might be sufficient to deform the epithelium through the holes in the smooth muscle mesh. The process of faveolar morphogenesis is relatively slow in these species (∼3–4 days in leopard gecko; ∼10 days in veiled chameleons), which unfortunately precludes experimental manipulation in culture. We therefore tested this hypothesis computationally using the finite element method. Specifically, we modeled the epithelium as a smooth, proliferative surface surrounded by a rigid lattice that matched the relative geometry of the squamate lungs (**Fig. 6A**). Increasing pressure of the lumen pushes the simulated epithelium through the holes in the smooth muscle mesh (**Fig. 6B**), as observed previously.^16^ Intriguingly, we found that in the absence of pressurization, growth of the simulated epithelium is also sufficient to expand this tissue through the mesh and generate a corrugated faveolar-like surface (**Fig. 6C**). Importantly, our computational model predicts that this proliferation-driven faveolar folding would also occur independently of epithelial thinning. To test this prediction experimentally, we measured the relative thickness of the epithelium during faveolar morphogenesis in all three species (**Fig. 6D–F**). As predicted, the faveolar epithelium of the brown anole was significantly thinner than that of both the leopard gecko and veiled chameleon (**Fig. 6G**), which were similar in thickness to each other. These data suggest that there are at least two physical mechanisms to fold the epithelium into faveolae: pressure-driven pushing and proliferation-driven expansion.

**Figure 6.**
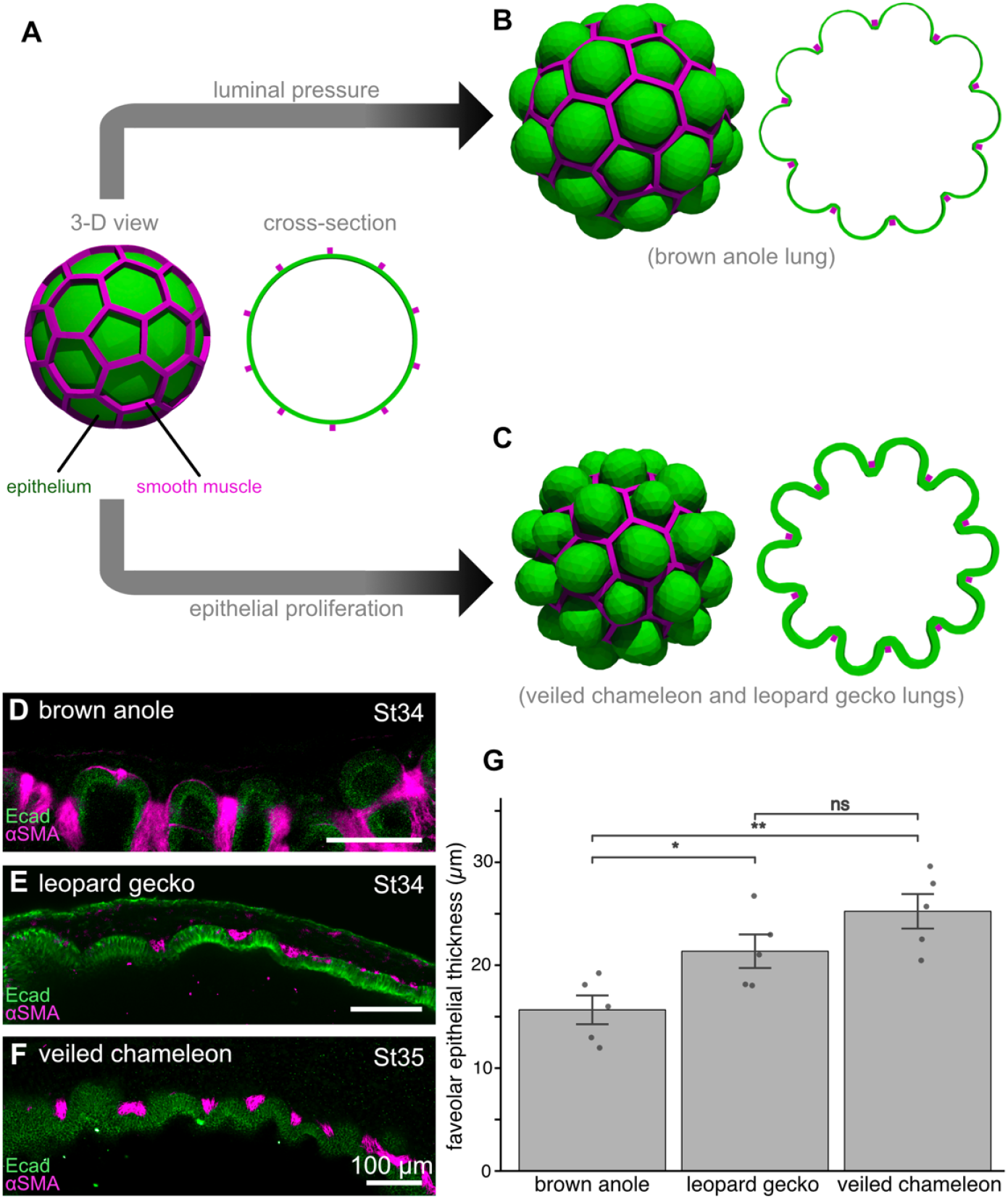
Computational simulations of epithelial deformations downstream of luminal pressure or epithelial growth predict epithelial thickness. **A**) The model begins as a smooth shell of epithelium (green) contained within a smooth muscle mesh (magenta.) The model experiences either increased luminal pressure (**B**) or epithelial growth (**C**). **D–F**) Z-slice images showing immunofluorescence for E-cadherin (green) and aSMA (magenta) in the developing faveolae of the brown anole (**D**), leopard gecko (**E**), and veiled chameleon (**F**). Scale bars, 100 µm. **G**) Graph showing epithelial thickness during faveolar morphogenesis. Asterisks and ns denote significant and non-significant differences, respectively, in pairwise *t*-test. Brown anole vs leopard gecko (*p*=0.029), brown anole vs veiled chameleon (*p*=0.003), leopard gecko vs veiled chameleon (*p*=0.14).

## 3 DISCUSSION

The lungs of brown anoles, leopard geckos, and veiled chameleons all start as wishbone-shaped epithelial tubes devoid of pulmonary smooth muscle. Around late stage 33, smooth muscle differentiates and subsequently organizes into a mesh. Based on our taxon sampling, differentiation of smooth muscle and remodeling into a mesh appear to be conserved across squamates. Consistent with our hypothesis, it was recently reported that the epithelium folds through holes lined by bundles of smooth muscle in the lung of the corn snake (*Pantherophis guttatus*).^36^ It remains unclear, however, what promotes pulmonary smooth muscle differentiation in squamates. Shh signaling is required for pulmonary smooth muscle differentiation in the murine embryonic lung.^8,9^ Similarly, blocking the Shh pathway with cyclopamine prevents smooth muscle differentiation in brown anoles, while activating Shh signaling with smoothened agonist results in ectopic smooth muscle.^16^ Additional work is needed to characterize the signaling pathways that regulate smooth muscle differentiation in squamates, broadly.

The subbronchi of chameleon lungs branch off of the main bronchus and elongate. After subbronchial elongation and faveolar morphogenesis, the diverticulae develop. These structures form in the absence of an obvious smooth muscle template, but show elevated pMLC at the apical surface of the epithelium, suggesting they may initiate via apical constriction, which folds the epithelium of the bird lung.^11^ Our results suggest that apical constriction might be conserved across avian and non-avian reptiles as a mechanism for pulmonary outgrowth and elongation. The molecular signals that induce the formation of subbronchi and diverticulae remain to be uncovered.

Fluid pressure serves as the physical force responsible for deforming the epithelium into faveolae in the anole lung.^16^ This inflation-driven process does not appear to occur during faveolar morphogenesis in either the leopard gecko or veiled chameleon. Instead, the actively growing epithelium appears to push itself through the holes in the smooth muscle mesh. This difference in physical mechanism may be the result of developmental timing. Brown anoles develop more rapidly than leopard geckos and veiled chameleons. Increasing luminal pressure in the anole lung generates faveolae quickly, instead of the multi-day proliferative process observed in geckos and chameleons. Consistent with this hypothesis, opportunistic embryonic sampling of the lacertoid Italian wall lizard (*Podarcis siculus*) revealed a decrease in aspect ratio during the period in which faveolae form (**Supplemental Fig. 1**), suggesting pressure-driven expansion in this faster developing species. It is also possible that developmental mechanisms are related to pulmonary physiology and/or specialization in the adult. Sampling additional squamate species^37^, including those that exhibit diversity of physiology in the adult and developmental lengths *in ovo*, may help to uncover additional developmental and mechanical constraints on epithelial morphogenesis and morphological diversification that enable ecological radiation and adaptation.

## 4 EXPERIMENTAL PROCEDURES

### 4.1 Lizard husbandry, egg collection, and dissection

We obtained eggs from a colony of wild-caught brown anoles (*Anolis sagrei*) housed at Princeton University (Princeton, New Jersey, USA). We performed husbandry, breeding, and maintenance as previously described^16^ and all animal procedures were approved by the Princeton University Institutional Animal Care and Use Committee (IACUC; protocol 2104). Briefly, we established ratios of one male to three females per enclosure, maintained at ambient temperature (23.9–29.4ºC) and humidity (>80%). We obtained eggs from a colony of captive-born leopard geckos (*Eublepharis macularius*) housed at Marquette University (Milwaukee, Wisconsin, USA). We performed husbandry, breeding, and maintenance as previously described^38^ and all animal procedures were approved by the Marquette University IACUC (protocols MU-4192 and MU-4241). Briefly, we established ratios of either one male to one female or one male to two females per enclosure. Further, we maintained ambient temperature (24.0–27.8ºC), a localized “warm spot” in each enclosure (28–31ºC), and ambient humidity (>75%). We obtained eggs from a colony of captive-born veiled chameleons (*Chamaeleo calyptratus*) housed at the Stowers Institute for Medical Research (Kansas City, Missouri, USA). We performed husbandry, breeding, and maintenance as previously described^32,39,40^ and all animal procedures were approved by the Stowers Institute for Medical Research IACUC (protocol 2023-160). Briefly, we housed chameleons individually and introduced females to male enclosures for <8 hours for breeding, maintained at ambient temperature (21–24ºC) and humidity (∼50%). Finally, we opportunistically obtained eggs from a colony of wild-caught Italian wall lizards (*Podarcis siculus*) housed at Princeton University. We performed husbandry, breeding, and maintenance similarly to brown anoles^16^ and all animal procedures were approved by the Princeton University IACUC (protocols 3154 and 3155).

Prior to dissection, we incubated eggs of all four species at Princeton University in a Vivarium Electronics VE-200 incubator in damp vermiculite at 26.7ºC. We removed embryos by submerging eggs in phosphate-buffered saline (PBS), piercing the eggshell using #5 watchmaker’s forceps (Fine Science Tools), and gently removing eggshell and extraembryonic membranes.^41,42^ We staged embryos using morphological criteria^30,31,33–35^ and selected embryos from stages 32 to 38. We dissected out lizard lungs by making three cuts using watchmaker’s forceps: one postcranial cut, one anterior to the pelvis, and finally removing the dorsal body wall and spine (**Supplemental Fig. 2**). After removing the dorsum of the embryo, we used forceps to gently remove the viscera and adjacent organs until we isolated the lung and trachea (**Supplemental Fig. 2**).

### 4.2 Immunofluorescence

We fixed lungs in 4% paraformaldehyde in PBS for 15–30 min at room temperature. We then washed samples four times in 0.1% Triton X-100 (Sigma-Aldrich) in tris-buffered saline (TBST) for 30 min each and then blocked for 1 hour at room temperature in a solution of 5% goat serum (Sigma-Aldrich) and 0.1% bovine serum albumin (Sigma-Aldrich) in 0.1% TBST. We placed samples in solutions of primary antibody targeting epithelial cytokeratins (CK; 1:100; DSHB, TROMA-I; 1:400; Dako, Z0622), E-cadherin (Ecad; 1:200; Invitrogen, 13-1900), α-smooth muscle actin (αSMA; 1:200; Abcam, ab5694; 1:400, Sigma-Aldrich, 1A4), and/or phosphorylated myosin light chain 2 (pMLC; 1:50; Cell Signaling Technology, 3675S) for 48 hours at 4°C. We then washed samples in TBST four times for 30 min each and subsequently placed samples in a solution of secondary antibody (1:400; Invitrogen) for 48 hours at 4ºC. We then dehydrated samples with serial dilutions of either isopropanol or methanol in PBS (25%, 50%, 75%, and 100%). Finally, we cleared samples using Murray’s clear (1:2 benzyl alcohol to benzyl benzoate; Sigma-Aldrich) and mounted samples in imaging chambers composed of nylon washers affixed to glass coverslips and filled with Murray’s clear.

### 4.3 EdU analysis

We visualized cell proliferation using a fluorescent assay for the thymidine analogue 5-ethynyl-2’deoxyuridine (EdU; Invitrogen C10638). We placed dissected lung explants on a semipermeable membrane (8-µm pore; Whatman) floating on Dulbecco’s modified Eagle’s medium (DMEM)/F-12 medium (Hyclone) supplemented with 5% fetal bovine serum (FBS; Gemini Bioproducts). We cultured samples at 37ºC for 30 min and then pulsed with EdU Click-iT reaction for an additional 30 min. Finally, we fixed lungs in 4% paraformaldehyde in PBS for 15 min at room temperature and proceeded to perform immunofluorescence in tandem with EdU (see 4.2).

### 4.4 Fluorescence imaging

We visualized fluorescently stained samples using either a Hamamatsu MAICO MEMS confocal unit fitted to an inverted microscope (Nikon Eclipse T*i*) or a Crest Optics X-Light V2tp confocal unit fitted to an inverted microscope (Nikon Eclipse T*i*2) with either 10× air or 20× air objectives. We acquired images using VisiView Software (Visitron Systems) with stage-stitching and z-series functions. We postprocessed images, adjusted image contrast and brightness, and made measurements using Fiji.^43^ We calculated lung aspect ratio for each species and developmental stage by measuring proximal-to-distal length and the medial-to-lateral width of the main bronchus (visualized using immunofluorescence for E-cadherin). We then divided the length by the width to obtain the aspect ratio.

### 4.5 *In silico* modeling of epithelial–smooth muscle interactions

We simulated the three-dimensional morphogenesis of a two-layered tissue structure using a custom finite element method (FEM) script developed in the Julia programming language, leveraging the Ferrite library.^44^ All simulation code and geometries are publicly available. This simulation follows the methods outlined previously.^16^ Briefly, we treated the processes of growth and pressure as much slower than mechanical relaxation.

Following finite deformation theory, we describe the mapping ***x*(*X*)** = ***X*** + ***u*(*X*)** from a material point ***X*** in the initial reference configuration *Ω*_0_ to a point ***x*** in the deformed configuration *Ω* via the displacement field ***u*(*X*)**. The local deformation is characterized by the deformation gradient tensor ***F*** = ***I*** + ***∇u***. To model growth, we decompose the deformation gradient as ***F*** = ***F***_***e***_***F***_***g***_. Here, ***F***_***g***_ is a tensor representing isotropic growth, defined as ***F***_***g***_ = *g****I***, where *g* is the growth factor and ***I*** is the identity tensor. The remaining deformation, 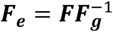, is the elastic deformation that generates mechanical stress.

The tissue layers are modeled as compressible neo-Hookean hyperelastic materials with a strain energy density function 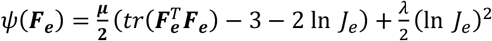, where *µ***(*X*)** = *E***(*X*)**/2(1 + ν**(*X*))** and *λ***(*X*)** = *E***(*X*)**ν**(*X*)**/(1 + ν**(*X*))(**1 − 2ν**(*X*))** are the Lamé elastic parameters for material at point ***X*** expressed in terms of the Young’s modulus *E***(*X*)** and Poisson’s ratio ν**(*X*)**, and ***J***_*e*_ = det**(*F***_*e*_) is the elastic volumetric deformation. This leads to the first Piola-Kirchhoff stress tensor

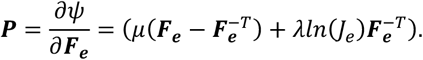

In mechanical equilibrium, the system satisfies equation *∇* · ***P*** = 0 in the bulk domain *Ω*_0_ with the boundary condition of the pressure *p* applied on the inner luminal surface of the epithelium *∂Ω*_*in*_ and no tractions applied on the external surface *∂Ω*_*out*_ · The displacement field ***u*** is then found by solving the weak form of the equilibrium for virtual displacements *δ****u***:

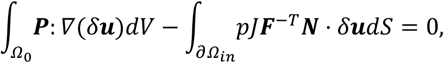

where we used integration by parts for the first term and used the Nanson’s formula for the second term to convert the virtual work of the pressure from the deformed configuration to the undeformed reference configuration, where ***N*** is the outward normal in the reference configuration. To ensure a unique solution, we constrained rigid body translation and rotation using Lagrange multipliers.

We discretized the governing nonlinear equations using first-order tetrahedral elements and solved incrementally using a Newton-Raphson iterative scheme. The simulation linearly ramped the growth factor *gg* and pressure *pp* to their final target values. The model used Young’s moduli of E=1.0 and E=10.0 for the epithelium and smooth muscle, respectively, with Poisson’s ratio of ν=0.3. Morphogenesis was driven by a total isotropic growth factor of g=2.0 within the epithelium, coupled with a final outward pressure of p=−0.2 applied to the lumen. Visualizations were generated in ParaView.^45^

## ACKNOWLEDGMENTS

We thank the laboratory animal facilities at Princeton, Marquette, and Rick Kupronis, Alex Muensch, and Marcus Wallas at the Stowers Institute for Medical Research for their support in live animal care.

Additional thanks to Aaron Bauer (Villanova University), who provided authors access to German dissertations on lizard lung anatomy.

## Abbreviations

αSMA: alpha smooth muscle actin
CK: cytokeratin
pMLC: phosphorylated myosin light chain

## Supplemental Figures

**Supplemental Figure 1.**
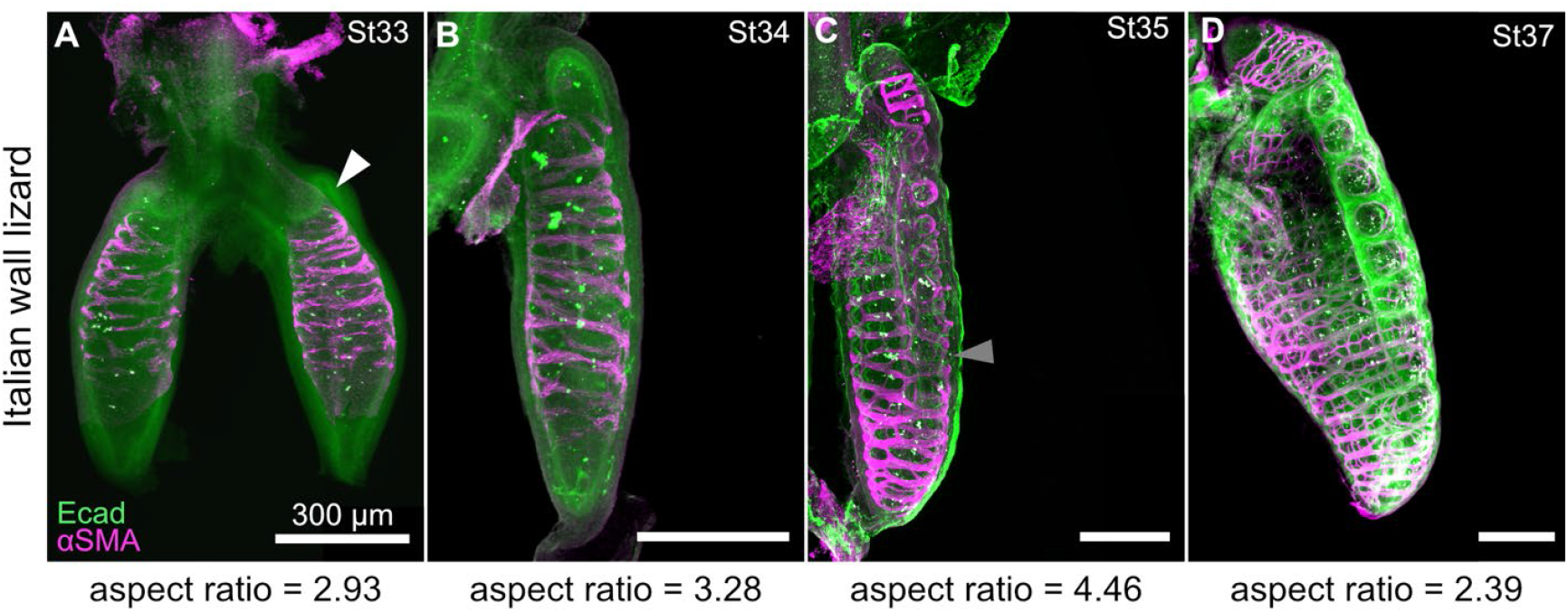
Opportunistic sampling, developmental staging, and aspect ratios of Italian wall lizard lungs. Images show immunofluorescence for E-cadherin and aSMA (**A**–**D**). St, developmental stage, Scale bars, 300 µm. White arrow denotes emergence of the proximal sub-chamber. Gray arrows denote epithelium pushing through the smooth muscle mesh.

**Supplemental Figure 2.**
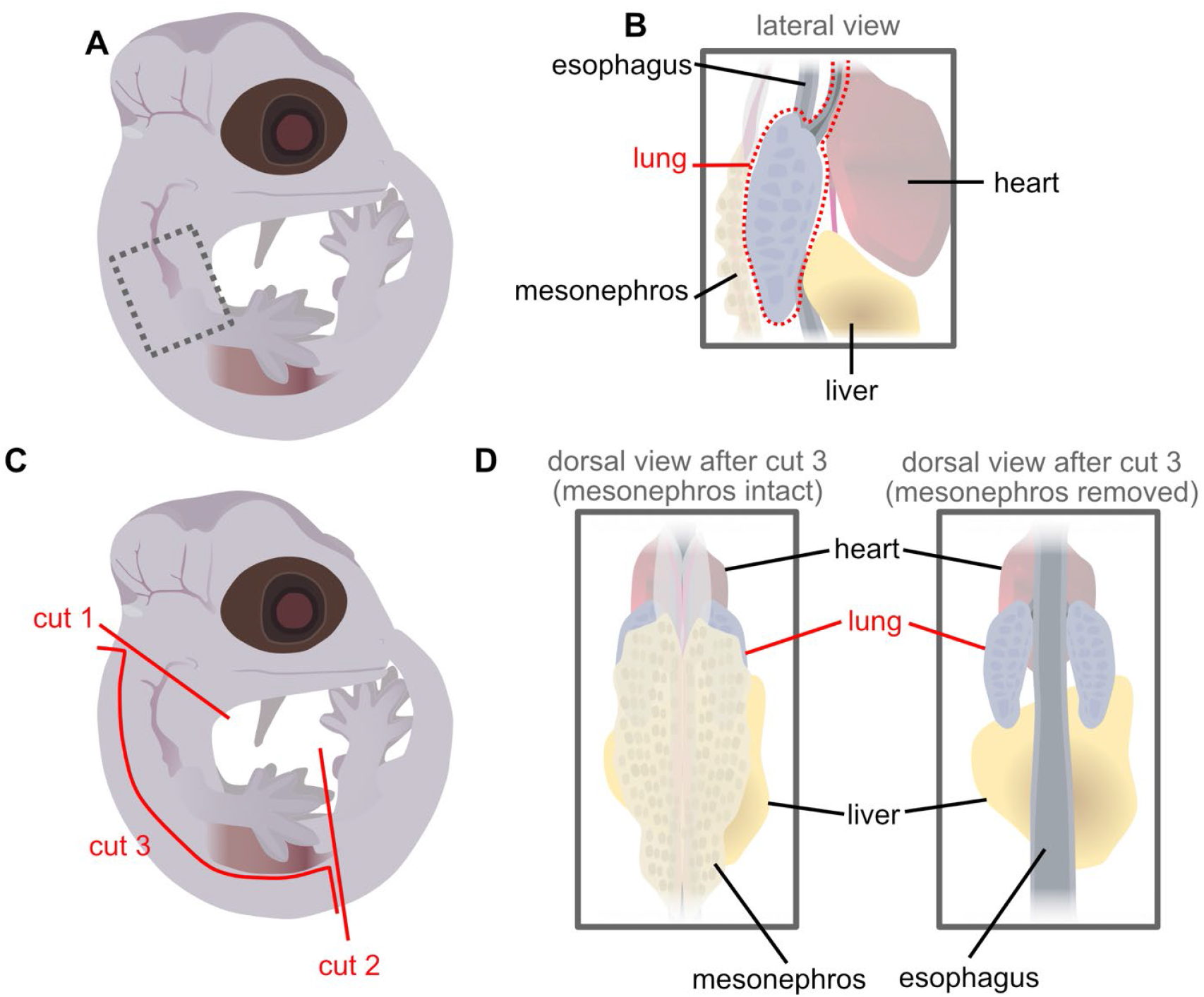
Schematic depicting procedure used to dissect lizard lungs. **A**) Illustration of a lizard embryo in lateral view. Dashed box corresponds to the region of the torso where lungs reside. **B**) Illustration of the lateral view of lizard viscera. **C**) Illustration of a lizard embryo in lateral view depicting regions to cut with dissection forceps (red lines). **D**) Illustrations of lizard viscera in dorsal view. The lung is visible after removal of the mesonephros.

## REFERENCES

1. Perry SF, Lambertz M, Schmitz A. Respiratory Biology of Animals. Oxford, UK: Oxford University Press; 2019.

2. Schachner ER, Hedrick BP, Richbourg HA, Hutchinson JR, Farmer CG. Anatomy, ontogeny, and evolution of the archosaurian respiratory system: A case study on Alligator mississippiensis and Struthio camelus. Journal of Anatomy. 2021;238:845–873.

3. Perry SF. Lungs: Comparative anatomy, functional morphology, and evolution. In: Gans C, Gaunt AS, eds. Biology of the Reptilia. Volume 19. Morphology G. Visceral Organs. Ithaca, New York, USA: Society for the Study of Amphibians and Reptiles; 1998:1–92.

4. Schachner ER, Cieri RL, Butler JP, Farmer CG. Unidirectional pulmonary airflow patterns in the savannah monitor lizard. Nature. 2014;506:367–370.

5. Schachner ER, Hutchinson JR, Farmer CG. Pulmonary anatomy of the Nile crocodile and the evolution of unidirectional airflow in Archosauria. PeerJ. 2013;1:e60.

6. Lambertz M, Grommes K, Kohlsdorf T, Perry SF. Lungs of the first amniotes: Why simple if they can be complex? Biology Letters. 2015;11:20140848.

7. Banavar SP, Fowler EW, Nelson CM. Biophysics of morphogenesis in the vertebrate lung. Current Topics in Developmental Biology. 2024;160:65–86. 10.1016/bs.ctdb.2024.05.003

8. Kim HY, Pang MF, Varner VD, Kojima L, Radisky DC, Nelson CM. Localized smooth muscle differentiation is essential for epithelial bifurcation during branching morphogenesis of the mammalian lung. Developmental Cell. 2015;34:719–726.

9. Goodwin K, Mao S, Guyomar T, et al. Smooth muscle differentiation shapes domain branches during mouse lung development. Development. 2019;146:dev181172.

10. Goodwin K, Lemma B, Zhang P, Boukind A, Nelson CM. Plasticity in airway smooth muscle differentiation during mouse lung development. Developmental Cell. 2023;58:338–347.

11. Kim HY, Varner VD, Nelson CM. Apical constriction initiates new bud formation during monopodial branching of the embryonic chicken lung. Development. 2013;140:3146–3155.

12. Tzou D, Spurlin JW, Pavlovich AL, Stewart CR, Gleghorn JP, Nelson CM. Morphogenesis and morphometric scaling of lung airway development follows phylogeny in chicken, quail, and duck embryos. EvoDevo. 2016;7:12.

13. Spurlin JW, Siedlik MJ, Nerger BA, et al. Mesenchymal proteases and tissue fluidity remodel the extracellular matrix during airway epithelial branching in the embryonic avian lung. Development. 2019;146:dev175257.

14. Palmer MA, Nelson CM. Fusion of airways during avian lung development constitutes a novel mechanism for the formation of continuous lumena in multicellular epithelia. Developmental Dynamics. 2020;249:1318–1333. 10.1002/dvdy.215

15. Rose CS, James B. Plasticity of lung development in the amphibian, Xenopus laevis. Biology Open. 2013;2:1324–1335.

16. Palmer MA, Nerger BA, Goodwin K, et al. Stress ball morphogenesis: How the lizard builds its lung. Science Advances. 2021;7:eabk0161.

17. Uetz P, Freed P, Aguilar R, Reyes F, Kudera J, Hošek J. The Reptile Database. Published online 2025. www.reptile-database.org

18. Klaver CJJ. Lung-morphology in the Chamaeleonidae (Sauria)and its bearing upon phylogeny, systematics and zoogeography. Journal of Zoological Systematics and Evolutionary Research. 1981;19:36–58.

19. Cieri RL, Craven BA, Schachner ER, Farmer CG. New insight into the evolution of the vertebrate respiratory system and the discovery of unidirectional airflow in iguana lungs. Proceedings of the National Academy of Sciences. 2014;111:17218–17223.

20. Kirschfeld U. Eine Bauplananalyse der Waranlunge. Zoologische Beiträge (Neue Folge). 1970;16:401–440.

21. Becker HO. Vergleichende Untersuchungen Am Respiratorischen Apparat Anguimorpher Eidechsen: Eine Stammesgeschichtliche Deutung. Dissertation. Rheinisch Friedrich-Wilhelms-Universität; 1993.

22. Wallach V. The lungs of snakes. In: Gans C, Gaunt AS, eds. Biology of the Reptilia. Volume 19. Morphology G. Visceral Organs. Ithaca, New York, USA: Society for the Study of Amphibians and Reptiles; 1998:93–296.

23. Werner F. Beiträge zur Anatomie einiger seltener Reptilien, mit besonderer Berücksichtigung der Atmungsorgane. Arbeiten aus dem Zoologischen Instituten der Universität Wien und der Zoologischen Station in Triest. 1912;19:373–424.

24. Bauer AM, Russell AP. A systematic review of the genus Uroplatus (Reptilia: Gekkonidae), with comments on its biology. Journal of Natural History. 1989;23:169–203.

25. Camp CL. Classification of the lizards. Bulletin of the American Museum of Natural History. 1923;48:289–481.

26. Perry SF, Duncker HR. Lung architecture, volume and static mechanics in five species of lizards. Respiration Physiology. 1978;34:61–81.

27. Perry SF, Duncker HR. Interrelationship of static mechanical factors and anatomical structures in lung evolution. Journal of Comparative Physiology. 1980;138:321–334.

28. Zheng Y, Wiens JJ. Combining phylogenomic and supermatrix approaches, and a time-calibrated phylogeny for squamate reptiles (lizards and snakes) based on 52 genes and 4162 species. Molecular Phylogenetics and Evolution. 2016;94:537–547.

29. Mauelshagen NMP. Die Phylogenetische Bedeutung Des Luftweg-Lungenkomplexes Der Gekkota. Dissertation. Universität Bonn; 1997.

30. Sanger TJ, Losos JB, Gibson-Brown JJ. A developmental staging series for the lizard genus Anolis: A new system for the integration of evolution, development, and ecology. Journal of Morphology. 2008;269:129–137.

31. Wise PAD, Vickaryous MK, Russell AP. An embryonic staging table for in ovo development of Eublepharis macularius, the leopard gecko. The Anatomical Record. 2009;292:1198–1212.

32. Diaz RE, Anderson CV, Baumann DP, et al. The veiled chameleon (Chamaeleo calyptratus Duméril and Duméril 1851): A model for studying reptile body plan development and evolution. Cold Spring Harbor Protocols. 2015;2015:889–894.

33. Dufaure JP, Hubert J. Table de développment du lézard viviparae (Lacerta vivipara Jacquin). Archives d’Anatomie Microscopique et de Morphologies Expérimentale. 1961;50:309–327.

34. Diaz RE, Shylo NA, Roellig D, Bronner M, Trainor PA. Filling in the phylogenetic gaps: Induction, migration, and differentiation of the neural crest cells in a squamate reptile, the veiled chameleon (Chamaeleo calyptratus). Developmental Dynamics. 2019;248:709–727.

35. Griffing AH, Sanger TJ, Daza JD, et al. Embryonic development of a parthenogenetic vertebrate, the mourning gecko (Lepidodactylus lugubris). Developmental Dynamics. 2019;248:1070–1090.

36. van Soldt BJ, Metscher BD, Richardson MK, Cardoso WV. Sox9 is associated with two distinct patterning events during snake lung morphogenesis. Developmental Biology. 2024;506:7–19.

37. Champini BG, Diaz RE, Schachner ER, Klein W. Pulmonary development in Squamata: Insights from embryonic studies using micro-CT. Developmental Dynamics. Published online In press. 10.1002/dvdy.70062

38. Thorogood J, Whimster IW. The maintenance and breeding of the leopard gecko, Eublepharis macularius, as a laboratory animal. International Zoo Yearbook. 1979;19:74–78.

39. Diaz RE, Anderson CV, Baumann DP, et al. Captive care, raising, and breeding of the veiled chameleon (Chamaeleo calyptratus). Cold Spring Harbor Protocols. 2015;2015:943–949.

40. Diaz RE, Bertocchini F, Trainor PA. Lifting the veil on reptile embryology: The veiled chameleon (Chamaeleo calyptratus) as a model system to study reptilian development. In: Sheng G, ed. Avian and Reptilian Developmental Biology. New York, New York, USA: Humana Press; 2017:269–284.

41. Sanger TJ, Hime PM, Johnson MA, Diani J, Losos JB. Laboratory protocols for husbandry and embryo collection of Anolis lizards. Herpetological Review. 2008;39:58–63.

42. Griffing AH, Sanger TJ, Matamoros IC, Nielsen SV, Gamble T. Protocols for husbandry and embryo collection of a parthenogenetic gecko, Lepidodactylus lugubris (Squamata: Gekkonidae). Herpetological Review. 2018;49:230–235.

43. Schindelin J, Arganda-Carreras I, Frise E, et al. Fiji: an open-source platform for biological-image analysis. Nature Methods. 2012;9:676–682.

44. Carlsson K, Ekre F. Ferrite.jl: A simple finite element toolbox written in Julia. Published online 2024. 10.5281/zenodo.13862652

45. Ayachit U. The Paraview Guide: A Parallel Visualization Application. Clifton Park, New York, USA: Kitware, Inc.; 2015.

